# Lycopene Production in Dedicated Novel Chasses for Lignocellulosic Waste Material Utilisation Capable of Sustained Coculture

**DOI:** 10.1101/2023.10.10.561056

**Authors:** John Allan, Matthew Crown, Matthew Bashton, Gary W Black

**Affiliations:** Department of Engineering Science, University of Oxford, Oxford, UK; Department of Applied Sciences, Health and Life Sciences, Northumbria University, Newcastle upon Tyne, UK

**Keywords:** municipal solid waste, synthetic biology, chassis organism, metabolic engineering, microbial communities

## Abstract

Municipal solid waste (MSW) represents tonnes of material that, for the most part, is relegated to landfill. Synthetic biology proposes solutions to many of the challenges faced by humanity today, but many approaches are confined to use in classical chassis organisms. In MSW there are a variety of potentially toxic materials such as glues, dyes, and preservatives that could pose a challenge to its capitalisation when using these commonplace chassis. We have isolated a bank of strains that utilise paper and cardboard waste from a relevant waste environment. From these we have identified three strains that are capable of utilising cellulose as a sole carbon source. We have analysed how they utilise cellulose and hemicelluloses, both alone and in coculture. This revealed insights to how they might be used in synthetic consortia which were then produced under laboratory conditions. Production of complete genome sequences of these strains provides genetic insight to how these processes may be occurring at the metabolic level, and how they could be augmented using synthetic biology. To this end, we have produced protocols for transforming plasmids into these strains and have produced high value metabolites from this material.

**Highlights:**

- Fully annotated genomes were produced from novel mesophilic aerobic strains isolated from lignocellulosic solid waste
- Lycopene was produced directly from relevant solid waste substrates by genetically modified variants of these strains
- Optimised carbon source blends influence coculture compositions of specific strains

## 1 Introduction

### 1.1 Paper waste and Lignocellulosic Biochemistry

Paper waste is composed of mainly lignocellulosic biomass. This is a rich seam of complex carbon which may be utilised by microbes. These complex carbon polymers are cellulose, hemicelluloses and lignin. Paper is manufactured using plant materials which in some cases can be recycled two to three times before their useful recyclable properties have expired. In others though, dyes, adhesives and other treatments to the material prevent it from being effectively recycled.

The complement of carbon-based polymers in paper materials could be rich seam of substrates and feedstock for biotechnology and consolidated bioprocessing. While a famously crystalline and recalcitrant material, many organisms have arisen which effectively utilise cellulose as a carbon source^1^. Hemicelluloses are a complex group of polysaccharides which crosslink cellulose fibres with lignin. They include xylan, polygalacturonans, arabinans, and hetero-polysaccharides like arabino-galactans. Once again, these form an effective carbon source for many microbes. The study of the enzymes which facilitate this has given rise to a field of research in glycobiology, and the functional super family of enzymes deserves its own classification as carbohydrate active enzymes (CAZymes)^2–4^.

Lignin has a random polyaromatic structure. The heterogeneity in the molecular architecture of the material has made it infamously challenging to degrade^5,6^. Nonetheless there are some groups of enzymes which attack it, *e.g*., laccases and peroxidases^7,8^. This polyaromatic structure however may be biotechnologically interesting as it reflects the structure of many industrially useful compounds, for example isoprenoids.

### 1.2 Chasses for Solid Waste Valorisation

Advances in synthetic biology and metabolic engineering have built effective pathways and platforms for the biosynthesis of various isoprenoids. These typically draw on central carbon metabolism, where there are two pathways which might be drawn upon to drive carbon flux towards isoprenoids, called the mevalonate^9^ and the more commonly found in bacteria DXP pathways^10–12^. The synthesis of carotenoid precursors to isoprenoids has been the focus of much metabolic engineering as these are a useful branchpoint to balance carbon flux in the pathway that is easy to measure and understand thanks to the ease of measuring these optically active intermediates. The carotenoid synthesis pathway utilised endogenously in many organisms involves three core genes called *crtE, crtB* and *crtI*. These encode enzymes wihch act in concert to produce lycopene from farnesyl pyrophosphate. This pathway has been discovered homologously in many microbes, insects and plants and has successfully been functionally reconstituted in *E. coli* and other common laboratory workhorses repeatedly^13–16^.

Recently a focus has emerged on the use of microbial communities in synthetic biology^17,18^. It is a current challenge to understand how best to support the requirements of many diverse strains simultaneously. Some work has investigated the use of complementary auxotrophic mutants of the same strain. In this scenario, each strain has near identical fitness where nutrients are replete, but each auxotroph may be selected for by the depletion of specific metabolites^19,20^. This may be sophisticated by the complementarity of auxotrophism, where strains may be reliant on the supply of specific metabolites from a partner strain. In other examples, strains may collaborate to tackle a single complex substrate, where each strain secretes non-redundant components of an enzymatic complex. Cybergenetics has also been deployed for long-term dynamic control, where a computational controller responds to fluorescent reporters indicative of specific strains, and responded with optogenetic regulation of antimicrobial resistance^21^. The use of diverse species within one of these synthetic communities has not been extensively explored to date.

It is the focus of many metabolic engineers to consider alternative feedstocks which are commonly considered as “waste” materials. This field of waste valorisation has made significant products^22^ from feedstocks that represent by-products of modern technologies and society. It makes sense to now turn our attention to utilising this lignocellulosic carbon “waste” in intelligent ways to produce specific high-value chemicals which might relieve our dependence on environmentally deleterious petrochemical sources.

While there are a variety of useful organisms for organisms for laboratory characterisation and tinkering, *e.g. E. coli, Saccharomyces cerevisiae… etc*., dedicated chasses may be of more use for bespoke functions. Various *Clostridia* have been adapted for utilisation of recalcitrant, yet abundant bagasse materials^23^. *Vibrio natrigens* is a remarkably fast organism for which genetic tools have been produced for quick DNA assembly pipelines, and circuit characterisation^24,25^. Production of glycosylated proteins and other eukaryotic natural products is facilitated by new tools in *Pichia pastoris*^26,27^. Carbon dioxide driven bioproduction might be achieved by turning to photoautotrophic microbes^28^ or even plants like *Marchantia polymorpha*^29^, or the easily-cultivated *Nicotiana bethamiana*^30^.

A dedicated suite of chasses for metabolic engineering utilising lignocellulosic material may then be of tantamount use to investigators interested in this feedstock. To this end we have sought to isolate, characterise and develop new strains from relevant paper sources.

## 2 Methods

### 2.1 Isolation and culture of MSW utilising bacteria

Samples of paper from an unsorted recycling waste bin were placed in a flask containing PCS medium. This medium has been used previously to enrich cellulose degrading organisms. After seven days, samples were plated on PCS agar plates (1.5% m/v agar, 0.5% m/v CMC). The colonies were then patched and the plates stained with Congo Red. Zones of clearing post staining indicated cellulase positive isolates. Cellulase positive isolates were cultured in LB medium and then stocked for future use.

Where defined carbon sources were utilised, the recipe JMM was utilised based on the recipe of Munoz *et al*^45^. An identical recipe was used to *Bifidobacterium* Media with the omission of resazurin and L-cysteine. EZ medium was obtained as a gift of Dr. Ciarán Kelly used as per the manufacturers instructions (Teknova). LB medium was used for routine culture.

For examinations of growth on different carbon sources the JMM medium was used and cells were cultured and measured autonomously using a Tecan Spark 10M platereader.

### 2.2 Flow cytometery

Cells were grown in the specified medium to exponential phase and a BD FacsCanto was utilised for flow cytometery. Wild type bacteria were used as a negative control to calibrate these measurements. The Flowing software package was used to analyse this data.

### 2.3 Electroporation

Plasmids were extracted by miniprep from *E. coli* DH5α hosts. Recipient strains were cultured in low salt LB up to an OD600 of approximately 0.3. The cells were then washed 4 times with ice cold water, and then concentrated 10-times in to 20% glycerol (m/v). Cells were electroporated with 100-200 ng of the relevant plasmid in 0.2cm gap BioRad electroporation cuvettes, at 2.5 kV, 200 ohms, and 25 uF. Cells were recovered with LB at 37 °C for approximately one hour and then plated on selective medium. 150 ug/mL kanamycin was required where appropriate and 25 ug/mL chloramphenicol.

### 2.4 Lycopene assays

Cells were cultured overnight in the relevant medium, according to the previous investigators^14,15,41^. Lycopene was quantified according to methods of Kim and Keasling^13^. Briefly, cells were harvested and resuspended in acetone and incubated at 55 °C for 15 minutes in the dark. The absorbance at 545 nm was then measured using a Tecan Spark 10M platereader. This was related to a series of standards of known concentration and normalised to OD600.

## 3 Results and Discussion

### 3.1 Isolation of cellulotrophic microbes

Many methods have been developed to produce enrichment cultures of microbes with specific phenotypes or other capabilities. The medium Peptone Cellulose Solution (PCS) has been utilised to produce communities of cellulose degrading organisms^31^. To enrich microbial strains which have adapted to a lifestyle on paper waste material, paper was sourced from an unsorted waste recycling bin. This was used to replace the cellulose in PCS medium as both a carbon source and inoculum.

These cultures were incubated statically for seven days at 37 °C. After this the cultures were plated on PCS agar plates and incubated overnight. The next day Congo red staining was used to identify cellulase positive strains and these were then held in strain libraries.

These isolates were then cultured in growth curves in JMM rich defined medium with the addition of CMC as a carbon source. The data was analysed using the GrowthcurveR package of software for R^32^ to identify the fastest growth rate (Supplementary Figure 1).

From this data, three roughly clustered groups could be identified. To avoid selection of strains isolated repeatedly, one strain from each of these groups was selected at random. By selecting isolates with distinct phenotypes in this was it was possible to ensure that three different strains were investigated. These three strains were US137-5, US137-6, US137-9.

To investigate deeper physiological phenotypes of these strains, they were investigated in growth curves identify their growth rates on a range of polysaccharides. CMC, polygalacturonic acid and xylan were investigated as carbon sources reflective of cellulose and hemicellulose. Additionally, simpler mono- and disaccharides were investigated which reflected the monomeric forms of these polysaccharides (Supplementary Figure 2). The hypothesis here being that an inability to utilise a polymeric carbon source may not mean an inability to use the sugar that it is formed of. Cross talk between these capabilities may also begin to reveal how these strains may interact. For this reason, cocultures were also examined, emulating the approach of Hardo *et al*^33^ where each strain was cocultured additionally with either one of the other two isolates. Increases in biomass yield upon coculture compared to monoculture may be explained by metabolic interactions between the strains. This revealed broad similarities in monocultures, for example monocultures had generally poor biomass yields when xylan was used as the carbon source. In cocultures however, where US137-6 was included an increased yield in biomass on xylan was observed.Genomic investigation and intervention Satisfied that these strains were useful assimilators of various relevant polysaccharides, and may be capable of synergistic interactions we moved on to sequence their genomes.

We were able to conclude that the strain US137-6 and US137-9 are distinct strains within the *Enterobacter cloacae* P101 complex by typing the 16S sequences using the Silva database^34^, with identities of 100% and 99.93% respectively. US137-5 was revealed as strain of *Klebsiella grimontii* using this also, with an identity of 99.86%, a close relative to *Klebsiella oxytoca* identified only in 2018^35,36^. None of these species have yet been deployed in contemporary synthetic biology studies to our knowledge. Hereafter these strains will be referred to as *E. cloacae* str. HBBE6, *E. cloacae* str. HBBE9, and *K. grimontii* str. HBBE5.

These strains contrast other studies of similar environments. Microbial ecologists have studied many environments where lignocellulosic material is digested. Bovine stomachs and bagasse fermenters typically appear to contain many thermophilic *Clostridia* among other organisms. In this work we have only genetically identified three strains, with these being strains of *Enterobacter cloacae P101*, and *Klebsiella grimontii*.. This is likely due to the culture-based approach here. Aiming to isolate microbes which are easy to grow in most laboratory environments, we grew microbes at 37 °C, and with no maintenance of anaerobic conditions. In this case then we would omit many of these thermophilic anaerobes.

Nonetheless, the three strains this study has focussed on are capable of digestion of many polysaccharides that compose paper waste. Indicating that functionally, mesophiles may be competent to degrade them. The mesophilic environment from which the inoculum was taken would also select for the maintenance of such organisms. While the paper may be destined for landfill of biomass digestion which would eventually select for the growth of these thermophilic anaerobes, it is possible that our enrichment of mesophilic aerobes represents a small study of the transient microbes that live on these materials while they fulfil their roles in the built environmentThe complete assemblies were annotated using the Bakta program^37^. Using this information, putative pathways were predicted using MinPath^38^, Figure 1A. This calculates the smallest number of pathways which can explain the putative functions in a set of genes. This analysis reveals a significant portion of each genome is dedicated to secondary metabolism. To investigate this further, the annotations were analysed using AntiSmash, which searches specifically for gene clusters which may be responsible for this and predicts what their functions may be. This reveals the presence of multiple clusters including those for aryl pigment synthesis, or polyketide synthases (Supplementary Material) *E. cloacae* str HBBE6 contains clusters responsible for putative pigment and aerobactin synthesis with high confidence. Clusters for RiPP-like antimicrobial products were found with highest confidence in the genome of *K. grimontii* str HBBE5. Expanded information on this can be found in Supplementary Data.

**Figure 1.**
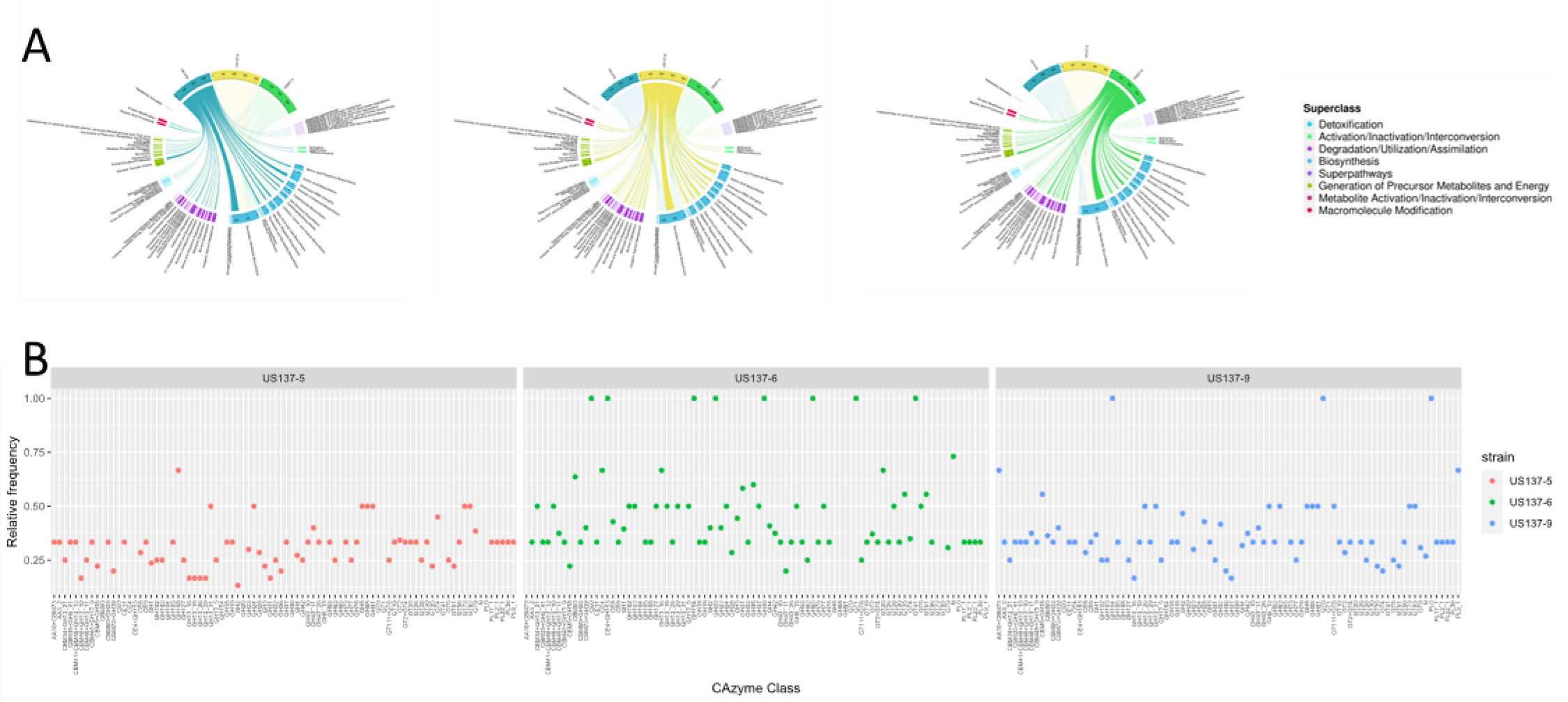
Each isolate selected encodes similar genomic characteristics, at a broad scale, with greater discrepancy in putative CAzymes. (A) Genomic DNA was isolated from each of the three selected isolates and sequenced using ONT and Illumina approaches. This was annotated using Bakta and the MetaCyc pathway classes predicted were used to produce Circos plots. (B) DBcan metaserver was used using these annotations to provide more specific predictions on CAZyme classes encoded. These are plotted here as the relative frequency of each enzyme class between strains.

The dbCAN metaserver for annotation of carbohydrate active enzymes^39^ was also utilised to annotate enzymes of this type with greater resolution. Frequencies of specific classes within each genome is shown in Figure 1B. *E. cloacae* str. HBBE6 shows the greatest number of unique classes of CAzyme, and *E. cloacae* str. HBBE9 the fewest. A number of unique enzymes between the three strains are found only in the strains *E. cloacae* str. HBBE6 and *K. grimontii* str. HBBE5.

DBcan annotation of these genome sequences displays a variety of different CAzyme types produced by each strain. Divergence in these annotation between each strain may explain some level of complementary activity we have observed in coculture experiments (Supplementary Figure 2). Both *E. cloacae* str. HBBE6 and HBBE9 appear to encode CAzymes of classes unique among these three genomes, with HBBE6 containing the most unique ones. Interestingly a class of LPMO is found in this genome which is not normally observed in *Enterobacter* genomes according to the CAZy database. This is a homolog of the *Vibrio cholerae* virulence factor GbpA, a chitin binding protein^42^. This enzyme should be studied further for its relevance to lignocellulosic waste material degradation, and whether or not its occurrence in this environment presents a novel function of this enzyme.

Chassis organisms must be readily transformable with new genetic material. To find if any regularly used plasmids might be transformed and maintained by strains US137-6, US137-9 and US137-5, electrocompetent preparations were made. They were then transformed with a variety of plasmids. ColE1, p15a and pBR322 origins proved to be suitable for plasmid maintenance (data not shown), with kanamycin selection at 150 μg/mL and chloramphenicol selection at 25 μg/mL where appropriate. The plasmid pGEO79-dasher, which encodes the dasherGFP allele under the control of a synthetic constitutive promoter and a ColE1 origin of replication, was obtained as a kind gift of Dr Paul James. This was transformed into each of these strains and single colonies cultured in LB medium or M9 or JMM media with the addition of 0.5% (m/v) CMC. The cultures were incubated overnight and then analysed by flow cytometry (Supplementary Figure 3). This provided insight to the homogeneity of expression between individual bacteria, as well as the degree of expression. The rich defined medium JMM provided the highest levels of dasherGFP expression in this case.

With these strains proven as being able to maintain synthetic DNA in the form of plasmids, we then set about investigating the potential of these strains as chasses for metabolic engineering. Isoprenoids are a high value group of chemicals which are often investigated by metabolic engineers as they can be produced from metabolites drawn from near universal pathways in central carbon metabolism. Since these chassis candidates are capable of assimilating various polysaccharides, we turned to see if the carbon assimilated in this way might be diverted to produce the isoprenpoid entry metabolite lycopene.

Genome sequences were utilised to predict metabolic pathways in these strains with the KAAS annotation server^40^ This predicts that these strains should be able to produce farnesyl pyrophosphate (FPP) from cellulose and other polysaccharides. This is strengthened by the observation of growth on this substrate also. Lycopene can be produced from FPP a pathway guided by the enzymes *crtEIB* in the DXP pathway. In order to test the hypothesis that lycopene can be produced in this manner by our candidate strains, we set about instigating their expression.

The plasmids pACCRT-EIB^14^ and pACHP-LYC^41^ were obtained as a kind gift of Prof Norihiko Misawa. pACCRT-EIB encodes the genes *crtE, crtB* and *crtI* from *Agrobacterium auranauticans*. pACHP-LYC encodes homologs of these genes from *Pantoea agglomerans* alongside the isopentyl phosphate isomerase *idi*. These were transformed into each strain along with the background empty vector pACYC184 as a negative control.

In the first instance single colonies were cultured in 2YT medium with the addition of 0.1 mM IPTG. Samples were taken from these cultures after overnight growth, and lycopene concentrations were estimated using the method of Kim and Keasling^13^ (Figure 2). Each plasmid produced statistically significant concentrations of lycopene compared to the negative control (p<0.005), however there was no significant difference between concentrations obtained between strains, under a Tukey-HSD test.

**Figure 2.**
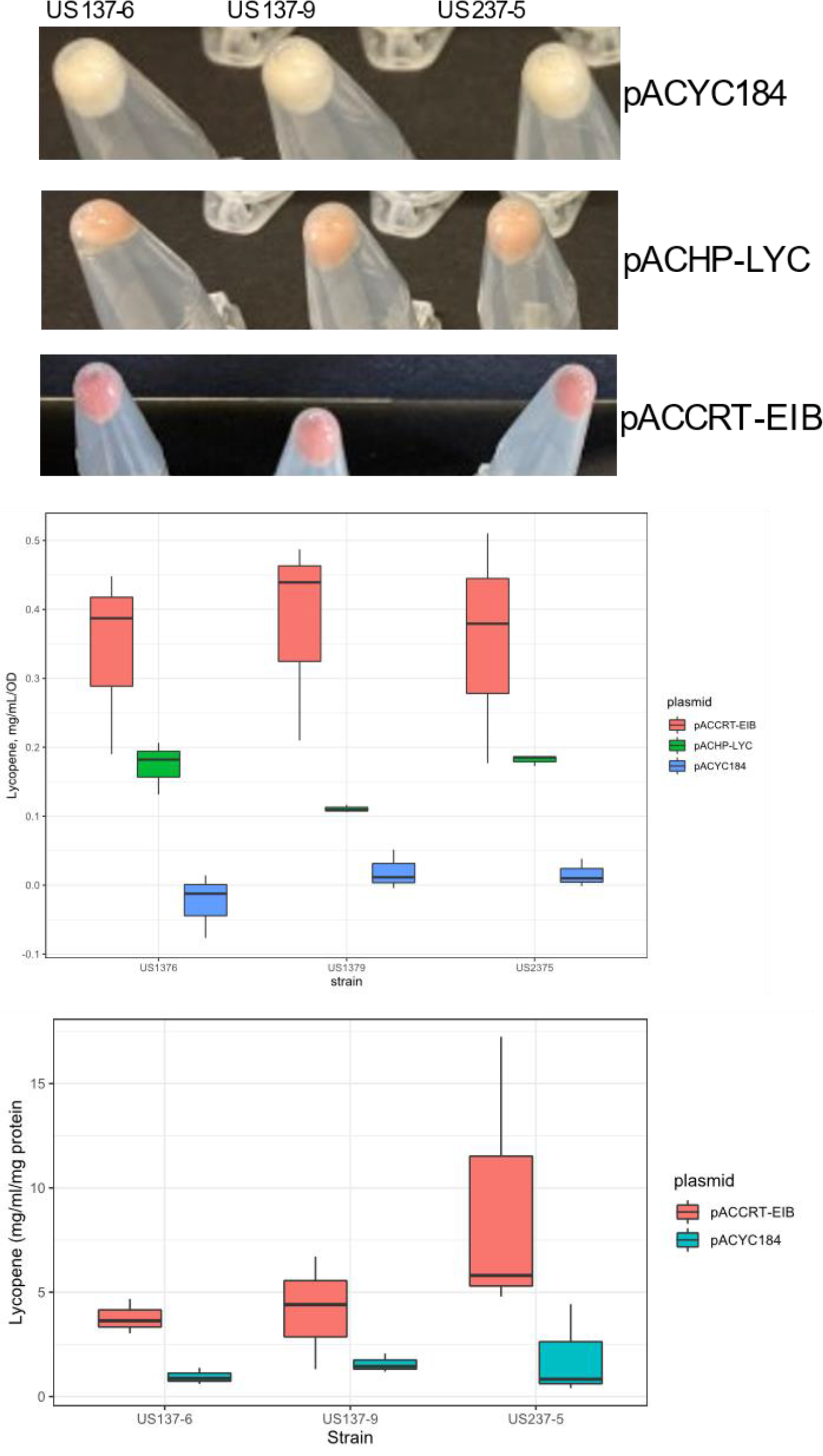
Lycopene can be produced directly from cellulose in each chassis candidate. (A) Chassis candidates were electroporated with the plasmids pACYC184, pACCRT-EIB, pACHP-LYC and cultured overnight at 37 °C in 2YT with the addition of 0.1 mM IPTG. Cultures were centrifuged and lycopene was measured by extraction with acetone and absorbance at 475 nm. (B) Measurements were normalised to the optical densities of the culture. (C) Strains harbouring the plasmids pACYC184 and pACCRT-EIB were cultured in EZ rich defined medium with 0.5 % (m/v) CMC as the sole carbon source. As in A, cells were harvested and lycopene measured. Concentrations were normalised to protein concentrations of the extracts.

Plasmid pACCRT-EIB instigated the highest concentrations of lycopene. This was taken forth to then study if lycopene could be synthesised directly from cellulose. In this case the rich defined medium EZ (Teknova) was used with the addition of 0.5% (m/v) CMC as a carbon source. This produced significant concentrations of lycopene in each strain where the pACCRT-EIB plasmid was used, however *K. grimontii* str. HBBE5 significantly outperformed the other two

Lycopene production is a common proof of concept in metabolic engineering. We have utilised that paradigm here to prove that these strains are capable of lycopene production from cellulose. Further experimentation should investigate other elements of central metabolism which may be tolerably drawn from for metabolic engineering, while also consider the use of other substrates. The abundance of lignin in lignocellulosic waste has not been explored in this work, however it presents potential as a precursor of many industrially relevant molecules. Future work could investigate the utility of dye-decolouring peroxidases^43^, laccases and other lignin specific enzymes in these strains. It may be useful to support central carbon metabolism in these scenarios with cellulose and hemicellulose, while strains degrade and reform lignin towards industrially interesting end-products.

The increased yield of biomass on specific substrates between strains presented the possibility to blend carbon sources to optimise the growth of two bacteria on a single carbon source. Utilising a full factorial screen of concentrations of CMC and xylan, the biomass yield on proportional blends of these carbon sources was investigated. Using two rounds of a design of experiments workflow we produced an surface estimation of the effect of this blend on biomass yield in cocultures of strains *E. cloacae* str. HBBE6 and *K. grimontii* str. HBBE5 (Figure 3A, Supplementary Figure 4).

**Figure 3.**
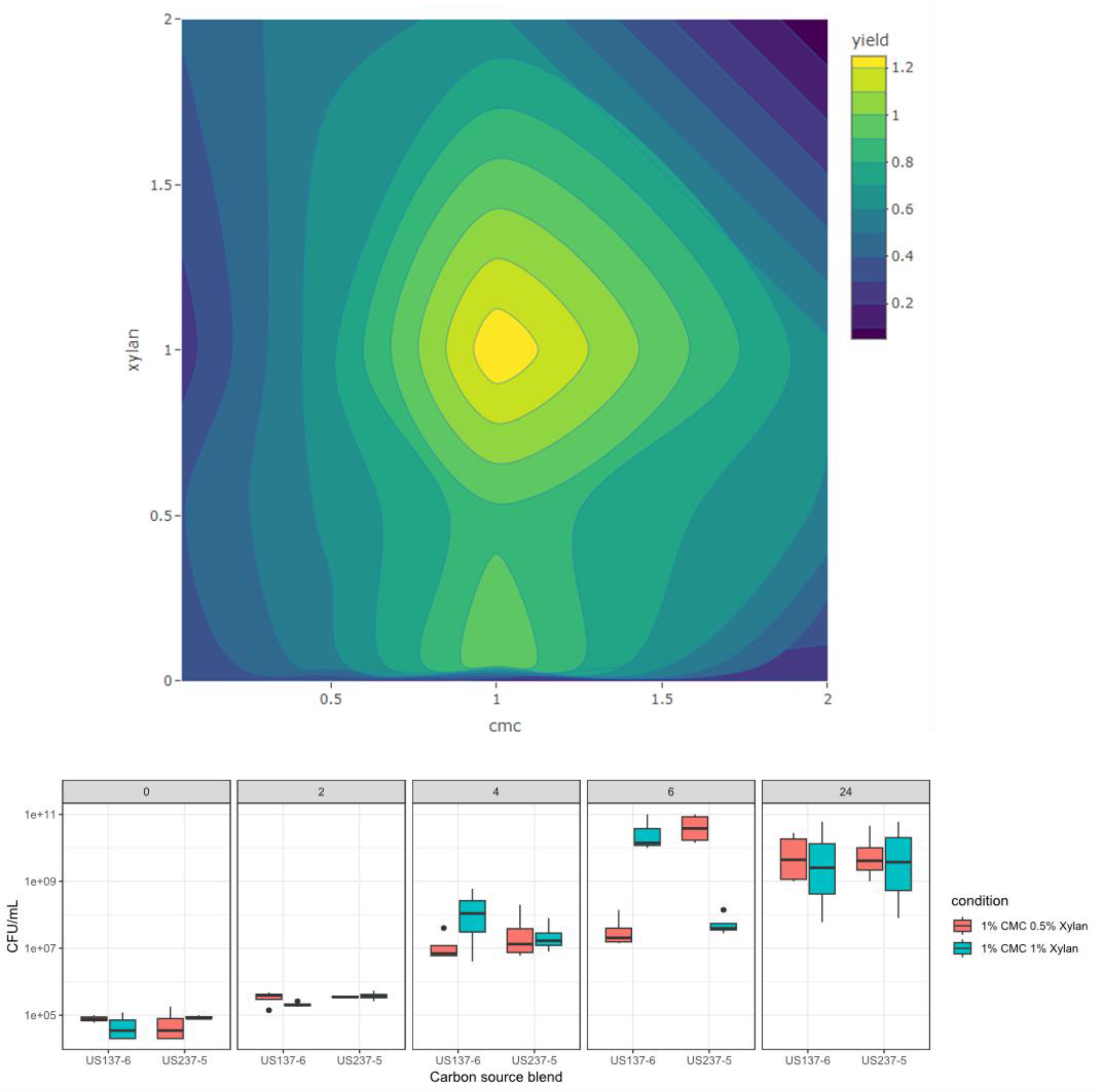
Design of Experiments based exploration of carbon source blends produces media compositions which can support divergent strains in coculture. (A) To find a blend of carbon sources which optimise the biomass yield of a coculture of strains HBBE5 and HBBE6, a full factorial design of experiments scoping and screening strategy was performed with concentrations of xylan and CMC between 0.05% and 1% (m/v). This was used to inform a second round of experiments with blends of the two polysaccharides between 0% and 2% (m/v). The data form both of these rounds of experimentation is displayed here. (B) Choosing the theorietically best blend of CMC and xylan from this data and a similar blend with less optimal predicted results, strains HBBE5 and HBBE6 were transformed with pGEO79dasher and pGEO79mOrange plasmids which encode dasherGFP and mOrange respectively. Samples were plated at 0, 2, 4, 6 and 24 hours post inoculation on selective agar and GFP positive and mOrange positive colonies were counted.

This predicted that concentrations of 1% (m/v) CMC and 1% (m/v) xylan would produce the greatest biomass yield, with a smaller peak ascending where xylan was reduced in concentration to nil. In order to investigate the dynamics of cultures grown in blends of carbon sources like these, the strains *E. cloacae* str. HBBE6 and *K. grimontii* str. HBBE5 were transformed with the plasmids pGEO79dasher and pGEO79mOrange respectively. The fluorescent reporters here permitted the tracking of each strain independently. At defined time points, samples were serially diluted and plated on selective media, with dasherGFP and mOrange producing colonies counted (Figure 3B). Six hours post inoculation, divergence between strain concentrations were observed, with *E. cloacae* str. HBBE6 being preferred in conditions with reduced xylan, and *K. grimontii* str. HBBE5 preferred in the higher xylan condition. This appeared to resolve at the later timepoint where the cultures have reached stationary phase, roughly equal concentrations of each strain were identified.

In future, the DOE approach could be used to develop more sophisticated blends of carbon sources and consider the inclusion of all three strains. It may also reveal ways to stabilise the community where genetic interventions may be made for metabolic engineering which disturb intrinsic interactions.

The observation that altering xylan concentrations produced differing compositions of strains at specific timepoints presents an interesting proposition for control of these microbial cocultures. The production and control of synthetic microbial consortia is a hotly researched topic in synthetic biology. We have demonstrated here a mechanism of supporting strains at roughly equal quantities, and at differing ratios depending on the concentration of specific carbon sources at specific cell densities. The Chi.bio^44^ platform permits turbidostat control of microbial cultures. This platform could be adapted relatively simply to control strain ratios on demand by maintaining a culture at a density which provides this strain concentration divergence, and altering the media composition at the user’s demand. In this way, a strain may be permitted to dominate the culture depending on which carbon source is supplied to the culture, leading to dynamic control of culture composition.

In future, carbon source blends should be investigated which support the culture of *E. cloacae* str HBBE9 in coculture. More interestingly, the production of useful metabolites in genetically modified strains should also be attempted in these cocultures.

By deploying genetic tools we were able to track the number of cells of each strain in coculture experiments. In all cases, both strains were culturable in comparable numbers to eachother by the end of the culture period. The coexistence of these bacteria, in contrived conditions even during stationary phase indicates competition may be reduced.

## 4 Conclusions

Here we have demonstrated that it is possible to move quickly from crude environmental isolation of microbes to their genetic modification and production of biotechnologically relevant metabolites. Our work here shows that chasses, which have typically been model organisms, can be rapidly developed and vindicated utilising a suite of contemporary tools and methods. Greater investigation will reveal how useful and robust they may be for a diversity of less contrived applications. We encourage their further development by interested colleagues.

## Supporting information

Genome analysis supplementary data

Main Body Figures

## 5 Acknowledgments

Dr. Paul James and Matthew Rogan should be thanked for their kind gifts of the pGEO79dasher and pGEOmOrange plasmids. Dr Ciarán Kelly and Emma Riley are thanked for the provision of EZ medium and Dr Kelly specifically for guidance in the development of this project. Prof Norihiko Misawa and Dr Miho Takemura are thanked for the gift of the plasmids pACYC184, PACCRT-EIB, and pACHP-Lyc. Dr Andrew Nelson, Dr Gregory Young and Prof Darren Smith should also be thanked for performing ONT sequencing for this study.

## 6 Author Contributions

JA and GB designed the study where JA performed all bench experimental work. JA assembled genomes while MC produced the annotations using BAKTA and MinPath which then are represented in Figure 1. MC was under the supervision of MB. The manuscript was compiled by JA and GB.

## 9 Supplementary Material

Supplementary Material should be uploaded separately on submission, if there are Supplementary Figures, please include the caption in the same file as the figure. Supplementary Material templates can be found in the Frontiers Word Templates file.

Please see the [Supplementary Material section of the Author guidelines](https://www.frontiersin.org/guidelines/author-guidelines#supplementary-material) for details on the different file types accepted.

## 12 Data Availability Statement

The datasets [GENERATED/ANALYZED] for this study can be found in the [NAME OF REPOSITORY] [LINK]. Please see the “Availability of data” section of [Materials and data policies in the Author guidelines](https://www.frontiersin.org/guidelines/policies-and-publication-ethics#materials-and-data-policies) for more details.

