## Supplementary figures and images for "Lycopene Production in Dedicated Novel Chasses for Lignocellulosic Waste Material Utilisation Capable of Sustained Coculture"

### s1 rates.png

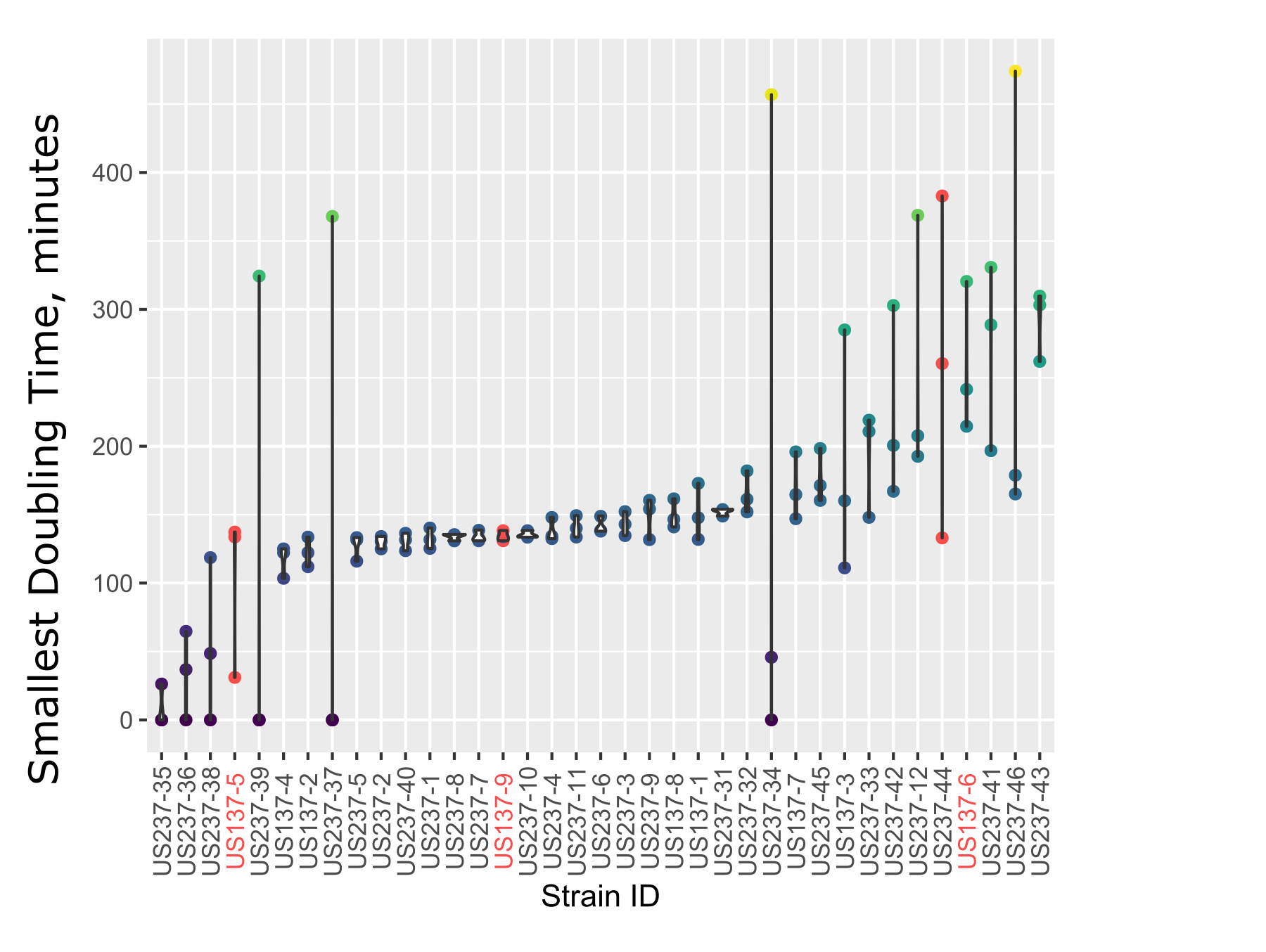
